# Shark sexing from forensic, archival, and developmental samples using sex-linked DNA markers

**DOI:** 10.64898/2026.05.02.722412

**Authors:** Akane Oishi, Yawako W. Kawaguchi, Taiki Niwa, Yoshinobu Uno, Shigehiro Kuraku

## Abstract

The effective management of threatened shark populations relies on accurate demographic data, particularly operational sex ratios. While sex identification in intact shark bodies is straightforward through the presence of external male organs, namely claspers, it remains impossible for processed fins in the illegal wildlife trade, early-stage embryos in breeding programs, or archived tissue fragments and blood samples where morphological traits are lost. Here, we present a robust molecular sexing framework leveraging recently identified sequences from shark sex chromosomes, consistently organized in the XY system, to our current knowledge. Our approach consists of two distinct methodologies tailored to the the current identification status of sex chromosome sequences in the target species. For the whale shark *Rhincodon typus* and the brownbanded bamboo shark *Chiloscyllium punctatum*, we employed end-point PCR assays targeting male-specific Y-linked markers. For the cloudy catshark *Scyliorhinus torazame*, we developed a quantitative PCR (qPCR) assay targeting differential X chromosome dosage. In this dosage-based system, females (XX) are distinguished by an amplification profile approximately one cycle earlier than males (XY). By integrating X-linked dosage quantification, our framework provides a critical internal control that significantly enhances reliability, allowing researchers to distinguish true females from PCR failures. This toolkit offers a versatile solution for diverse applications, ranging from the study of sex determination mechanisms in pre-phenotypic embryos to the reconstruction of sex ratios from space-constrained tissue archives and global wildlife forensics, thereby contributing to the comprehensive conservation of shark biodiversity.

## Introduction

Chondrichthyans (cartilaginous fishes) consisting of sharks, rays, and chimaeras, are among the most threatened vertebrates globally, with over one-third of species estimated to be at risk of extinction due to overfishing and habitat loss (Dulvy et al., 2021). Effective conservation management relies heavily on understanding population dynamics, particularly the operational sex ratio. Many chondrichthyan species exhibit sexual segregation, where males and females occupy different habitats or depths (Klimley, 1987; Sims et al., 2001; Norman & Stevens, 2007; Wearmouth & Sims, 2008; Mucientes et al., 2009; Laso-Jadart et al., 2025). Consequently, sex ratio imbalances within local populations inevitably lead to disproportionate catches of one sex. Establishing the ability to determine the genetic sex from the body parts of landed individuals enables sex-aware resource surveys that can provide a more accurate picture of natural populations; however, it presents a significant challenge when morphological identification is impossible.

While sex identification in intact chondrichthyan specimens is straightforward due to the presence of claspers in males (Gayford, 2023), this morphological trait is absent in dressed carcasses, processed fins found in the illegal wildlife trade, and early-stage embryos analyzed in developmental biology. In such cases, molecular sex identification, conventionally performed in mammals (Sullivan et al., 1993; Aasen & Medrano, 1990) and birds (Griffiths et al., 1998; Fridolfsson & Ellegren, 1999), becomes indispensable. In teleost fishes, sex-determining genes [e.g., *dmy* in medaka (Matsuda et al., 2002), *sdY* in salmonids (Yano et al., 2012; Yano et al., 2013)] and PCR-based markers are well-established (e.g., Zhu et al., 2024). In contrast, the sex-determination systems of cartilaginous fishes have long remained elusive, although recent genomic advances have begun to shed light on their sex chromosomes (Fig. 1; Yamaguchi et al., 2023; Sendell-Price et al., 2023; Niwa et al., 2025; Dahms et al., 2025; also see Uno et al., 2020 for a recent cytogenetic effort). So far, the application of sex-linked markers in sexing, which has become a general practice (e.g., Houchen et al., 2024), has not been routinely exercised in chondrichthyans.

**Figure 1.**
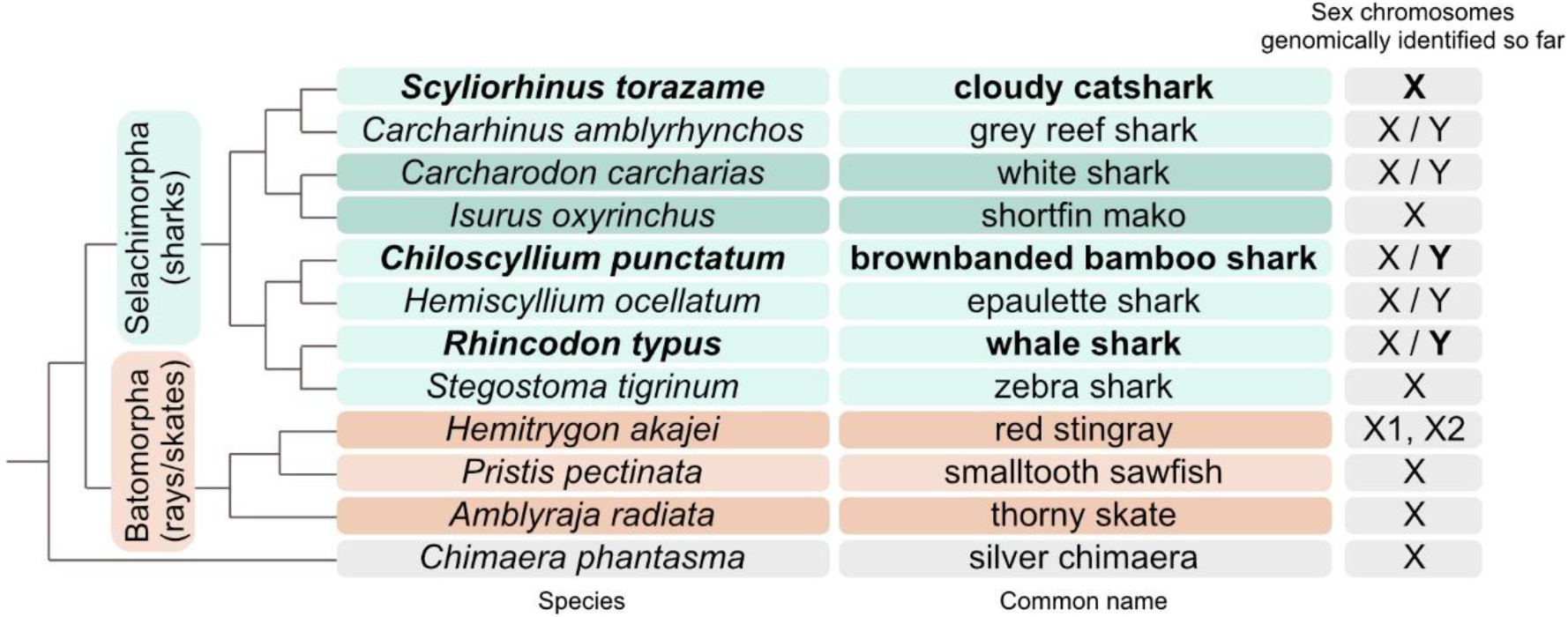
Taxonomic distribution of sex chromosome identification among chondrichthyans. This figure contains only the sex chromosomes identified through male-female comparisons of genome sequences (Yamaguchi et al., 2023; Sendell-Price et al., 2023; Niwa et al., 2025; Dahms et al., 2025; Teramura et al., 2026; Lee et al., 2025) and excludes cytogenetic observations. The species targeted in the present study as well as the sex chromosome targeted for our PCR experiments are shown in bold. Adjacent species in the same background color belong to the same taxonomic order. Among the species included in this figure, the silver chimaera, a holocephalan species, is an exception in that its X chromosome, identified as a fragment, has a distinct autosomal origin from elasmobranchs (Teramura et al., 2026).

In this study, we developed a robust molecular sexing method based on DNA sequence information and PCR amplification. We established this method using high-quality genomic resources from phylogenetically distinct species: threatened biggest fish species that is commercially relevant in some regions, the whale shark *Rhincodon typus*, as well as the brownbanded bamboo shark *Chiloscyllium punctatum* used recently for developmental biological studies, and the cloudy catshark *Scyliorhinus torazame*. This study provides a versatile and reliable toolkit for sex identification across diverse chondrichthyan lineages, facilitating better monitoring of illegal trade for wildlife forensics and early development before morphological sexual dimorphism appears.

## Methods

### Animals

For the whale shark *Rhincodon typus*, we used genomic DNA extracted for our earlier studies (Yamaguchi et al., 2023; Kawaguchi et al., 2026), from individuals kept at aquariums. We also used the samples of the brownbanded bamboo shark *Chiloscyllium punctatum* (BioSample SAMN49113356-SAMN49113360, SAMD00098917) and cloudy catshark *Scyliorhinus torazame* (SAMN49113389-SAMN49113392), both prepared originally for our previous study (Niwa et al., 2025). The details of the samples used in the present study are included in Supplementary Table 1. The animals were unambiguously sexed by the presence and absence of the claspers, male-specific structures on pelvic fins.

Animal handling and sample collections at the aquariums were conducted by veterinary staff without restraining the individuals, in accordance with the Husbandry Guidelines approved by the Ethics and Welfare Committee of Japanese Association of Zoos and Aquariums. All other experiments were conducted in accordance with the Guideline of the Institutional Animal Care and Use Committee (IACUC) of RIKEN Kobe Branch (Approval ID: H16-11), National Institute of Genetics (Approval ID: R5-14 and R6-13).

### DNA extraction

High-molecular-weight genomic DNA of whale shark, extracted previously for whole genome sequencing (Kawaguchi et al., 2026), was sheared to achieve high precision of reaction preparation with modest viscosity and retrieve a major DNA size distribution of 10-50 kb. For DNA fragmentation, 2 µg of genomic DNA was diluted to a final volume of 150 µL with TE buffer and loaded into a g-TUBE (Covaris). Centrifugation was performed at 25°C using a TOMY MX300 centrifuge equipped with a TOMY AR015-24 rotor. The sample was sequentially centrifuged at 6,100 rpm for 1 min, 6,100 rpm for 1 min, and 7,400 rpm for 1 min. The g-TUBE was then inverted and centrifuged again under the same conditions (6,100 rpm for 1 min, 6,100 rpm for 1 min, and 7,400 rpm for 1 min). For all other individuals including both brownbanded bamboo shark and cloudy catshark subjected to our previous study (Niwa et al., 2025), genomic DNA was extracted with Monarch Genomic DNA Purification Kit (New England Biolabs).

### Primer design

Oligonucleotide primers for both end-point PCR and quantitative PCR (qPCR) were designed using Primer3Plus (Untergasser et al., 2012). Target amplicon sizes were set to 80–200 bp for end-point PCR and 75–100 bp for qPCR. Candidate primer pairs were evaluated and selected based on low thermodynamic penalty scores. For targets within protein-coding genes, primers were strictly positioned within single exons to avoid intronic regions.

### End-point PCR

DNA fragments were amplified using ExTaq DNA polymerase (TaKaRa Bio) with oligonucleotide primers of 0.8 μM each and 35 ng of genomic DNA templates on a GeneAmp PCR System 9700 (Applied Biosystems), in the following thermal cycling conditions: initial denaturation for 3 min at 98 °C followed by 30 cycles of denaturing for 10 seconds at 98 °C, annealing for 30 seconds at 60 °C, and extension for 3 mins at 72 °C, after which the reaction tube was kept at 12 °C. The PCR products were loaded onto 3% agarose gel, together with 100 bp ladder marker (TaKaRa Bio).

### Quantitative real-time PCR

After expected amplification is ensured with end-point PCRs performed as above, quantitative PCRs were performed using Luna Universal qPCR master mix (New England Biolabs) with 0.25 μM of each oligonucleotide primer and 35 ng of genomic DNA templates on CFX96 Real-time PCR System (Bio-Rad), in the following thermal cycling conditions: initial denaturation for 1 min at 95 °C followed by 40 cycles of denaturing for 15 seconds at 95 °C and annealing and extension for 30 seconds at 60 °C, after which additional steps for dissociation curve analysis was performed with increment of 0.5 °C every 5s from 65 °C to 95 °C. All samples were run in triplicate. Amplification of a single specific PCR product was confirmed with melting curves. Quantification was performed with CFX Maestro Software (Bio-Rad). The R^2^ of the standard curve and the dissociation curve were also analyzed to confirm specific amplification and quantification of the target region. Relative copy numbers were calculated using the *Δ*Cq method, with autosomal markers serving as the baseline for determining the double-dosage expected in females.

### Results

### Sexing for whale shark based on male-specific markers on Y chromosome

Previous whale shark genome analysis revealed genomic DNA sequences that showed male specificity, identified as putative Y chromosome fragments (scaffold0112 [JAHMAH020000112.1] and scaffold0679 [JAHMAH020000679.1]; Kawaguchi et al., 2026). These male-specific sequences are scarce in protein-coding genes and harbor massive repetitive sequences, which we avoided in designing two pairs of PCR primers as a male marker in the present study (Fig. 2B; Table 1). In the present study, we designed a male-specific primer set on one of the previously identified Y chromosome fragment (scaffold0112, positions 5406 to 6100). As a control, we also designed a PCR primer pair for *HoxA1* gene located in an autosome (chromosome 2).

Using these PCR primer sets, we performed genomic DNA amplification for two male and female individuals, respectively, for which sex is unambiguously determined morphologically (see Methods) (Fig. 2C).

**Table 1.**
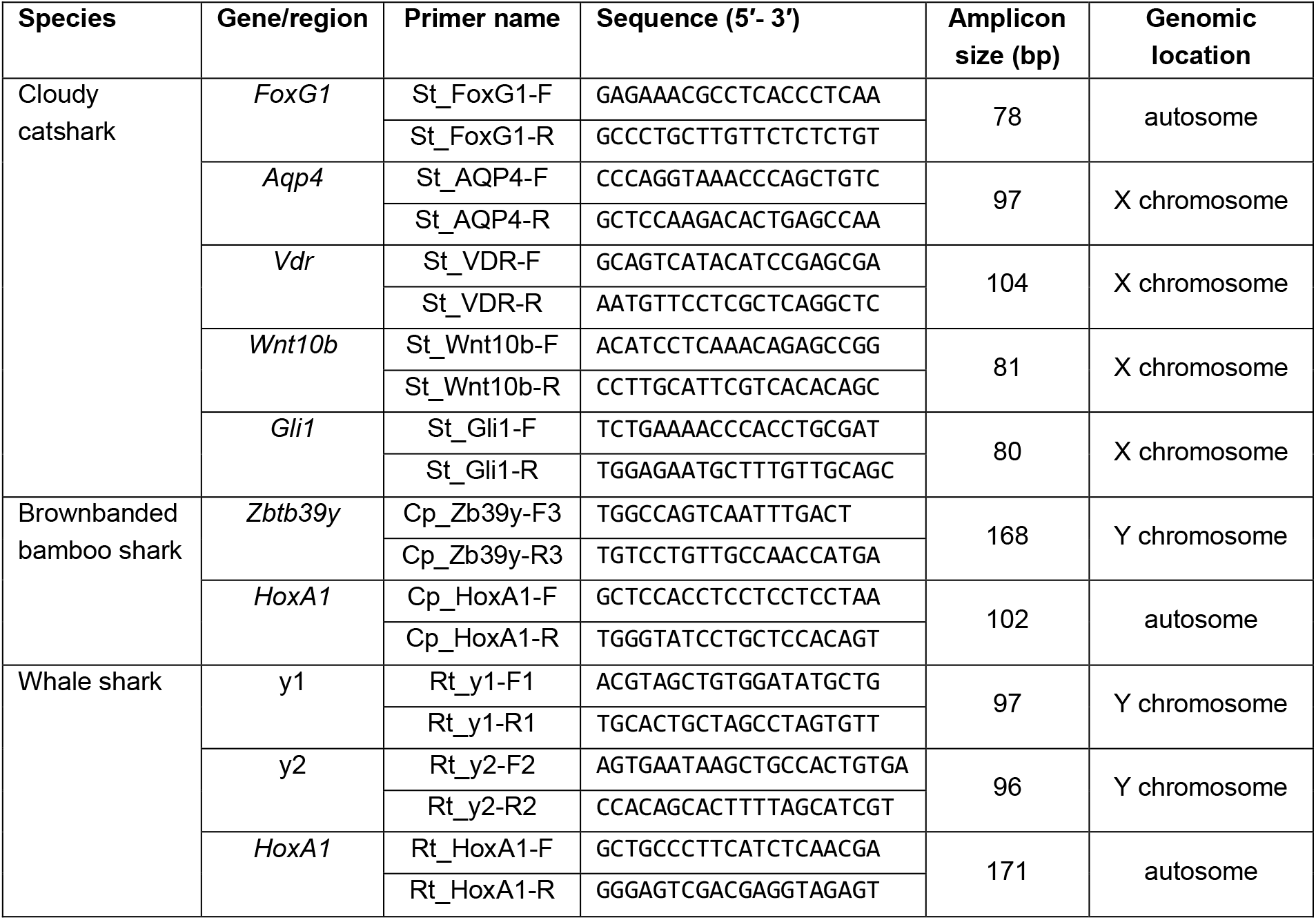
Primers designed to amplify sex-linked and autosomal markers in the present study.

**Figure 2.**
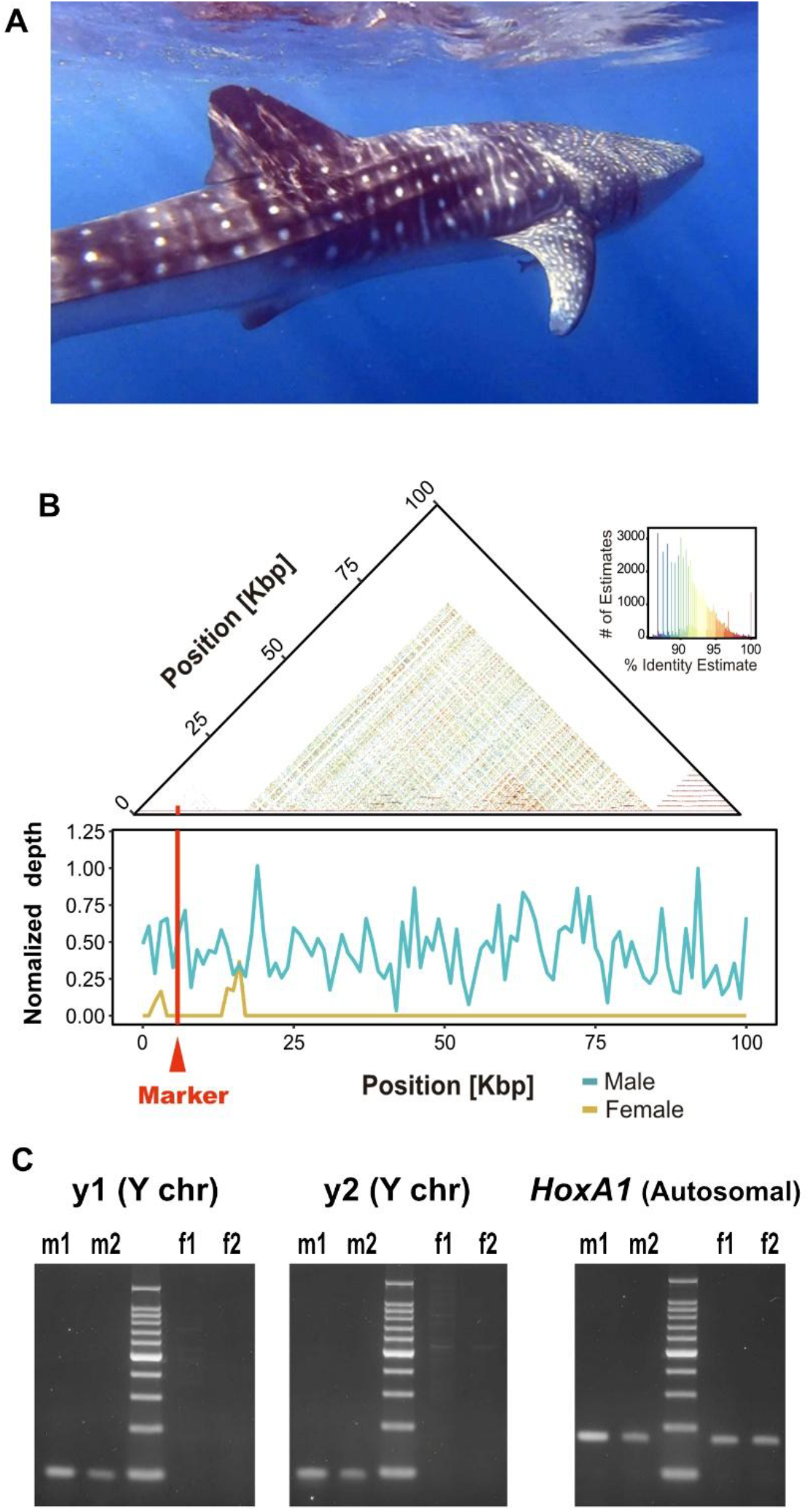
Sexing for whale shark. A, Whale shark. Photo credit: Shigehiro Kuraku. B, Designing PCR primers on the whale shark Y chromosome sequences. Self-identity dot plot generated with ModDotplot (Sweeten et al., 2024) and normalized read-depth profiles of the putative Y chromosome fragment scaffold0112 in short-read whole genome sequencing (Kawaguchi et al., 2026). The PCR marker, indicated by the red vertical line (‘Marker’), was chosen in a region with weak tandem-repeat signals and unambiguous male-specific coverage. Normalized depth represents read depth averaged in 1-kb windows and divided by the genome-wide mean depth. C, Electrophoresis of PCR products on 3% agarose gel. Amplification was tested using two Y-chromosome marker primer sets, y1 (Amplicon size: 97 bp) and y2 (96 bp), alongside the autosomal control *HoxA1* (171 bp). Templates included gDNA from two males (m1 and m2) and two females (f1 and f2). A 100-bp DNA ladder was used as a size marker.

Consistently, we observed positive PCR bands for males, but not for females, for both Y chromosome markers, while both males and females allowed amplification for the autosome marker. The amplification of the autosomal marker for females guarantees the valid PCR condition, which cannot be explained by any failure in DNA extraction or PCR amplification.

### Sexing for brownbanded bamboo shark targeting Y chromosome

Previous genome analysis for brownbanded bamboo shark (Fig. 3A) revealed genomic DNA sequences that showed male specificity, identified as a putative Y chromosome (Niwa et al., 2025; NCBI, NC_092792.1). The Y chromosome sequences are scarce in protein-coding genes with massive repetitive sequences. In the present study, we avoided the repetitive regions and designed a pair of PCR primers as a male marker in the protein-coding exon of the *Zbtb39* ortholog (Figure 3B of Niwa et al., 2025; Table 1). Because this gene has two genomic copies with few nucleotide differences (X type [NCBI Gene ID:140471229] and Y type [NCBI Gene ID:140471380]), we selected the genomic segment in which only the Y type (*Zbtb39y*) is amplified using a specific primer set (Fig. 3B). As a control, we also designed a PCR primer pair for *HoxA1* gene located on an autosome (chromosome 8; Kuraku, 2026).

**Figure 3.**
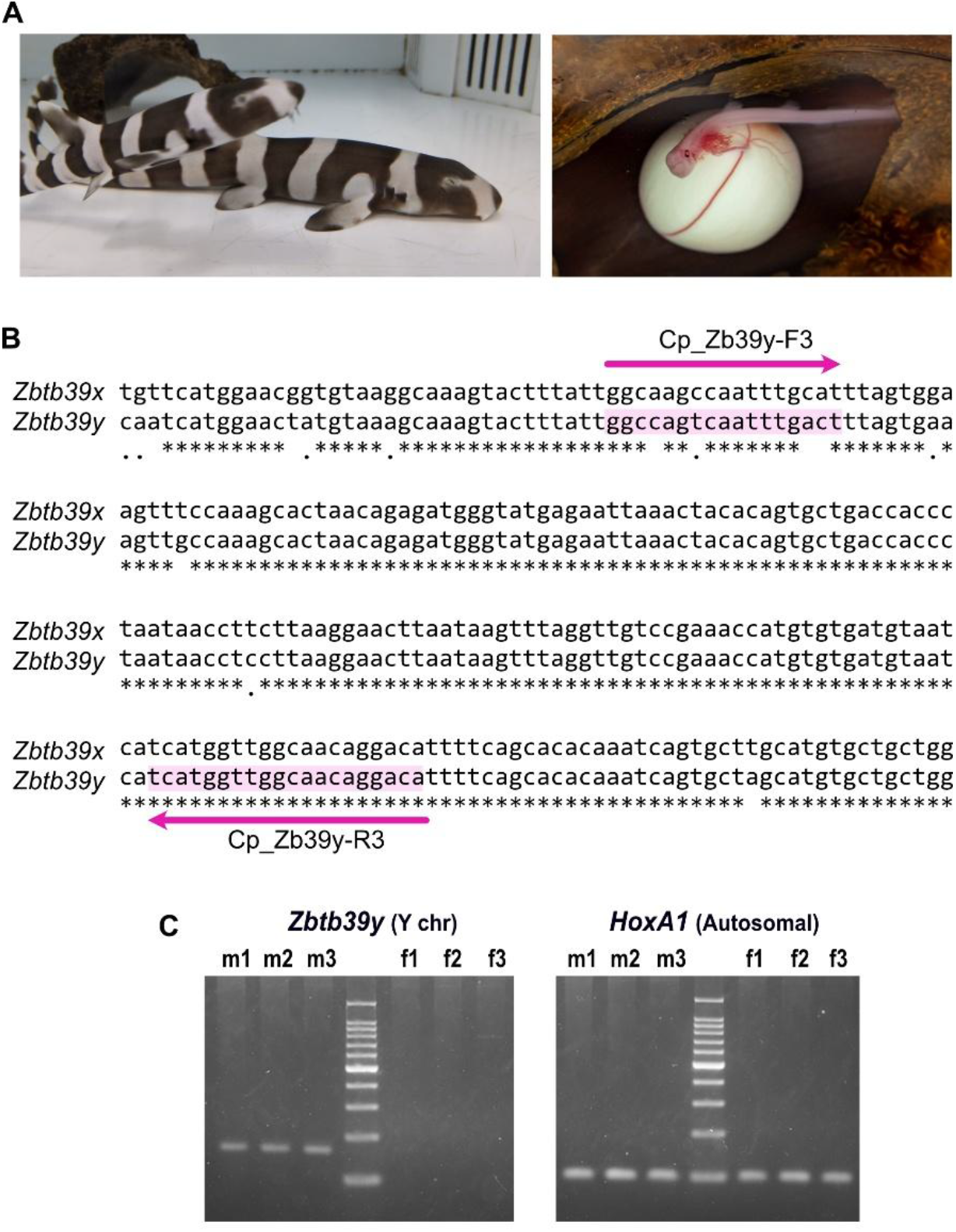
Sexing for brownbanded bamboo shark. A, Brownbanded bamboo shark juveniles (left) and developing embryo in the windowed eggcase (right). Photo credit: Shigehiro Kuraku. B, Location of the designed primers. Nucleotide sequences are shown for the region starting at position 198 of the *Zbtb39y* transcript (XM_072567404.1). The forward primer harbors several mismatches between the X-type and Y-type sequences previously identified (Niwa et al., 2025), allowing no amplification in females with the adopted PCR condition. This genomic region harbors no introns. C, Electrophoresis of PCR products on 3% agarose gel. Amplification was performed using primer sets targeting the Y-linked *Zbtb39* gene (expected amplicon size: 168 bp) and the autosomal *HoxA1* gene as a positive control (102 bp). Genomic DNA from three males (m1–m3) and three females (f1–f3) was used as templates. A 100-bp DNA ladder was used as a size marker.

Using these PCR primers, we performed genomic DNA amplification in triplicates for three male and female individuals, respectively, for which sex is unambiguously determined morphologically (see Methods; Fig. 3C). Consistently, we observed positive PCR bands for males, but not for females, for the Y chromosome markers, while both males and females allowed amplification for the autosome marker *HoxA1*.

### Sexing for cloudy catshark based on X chromosome dosage

To date, no Y chromosome sequence has been identified in catsharks (the genus *Scyliorhinus*; Fig. 4A) (Mayeur et al., 2024; Niwa et al., 2025), but our previous comparative elasmobranch genome analysis revealed the X chromosome of the cloudy catshark, possessed in double dosage in females (Niwa et al. 2025). Despite the limited knowledge, we focused on the differential among-sex dosage of X chromosome for sex identification. To design PCR primer pairs, we chose single orthologs of four protein-coding genes (*aquaporin 4 [Aqp4*, see Kuraku et al., 2024], *vitamin D receptor* [*Vdr* or *NR1I1*], *Wnt10b* [see Kuraku 2021], and *Gli family zinc finger 1* [*Gli1*]) on the 46 Mb-long X chromosome (NCBI, NC_092738.1; Niwa et al., 2025) that showed a comparable gene density to autosomes (Table 1). As a control, we also designed a PCR primer pair for *FoxG1* gene (Hara et al., 2018) located in an autosome (chromosome 2) for which we expect equal dosage between males and females.

**Figure 4.**
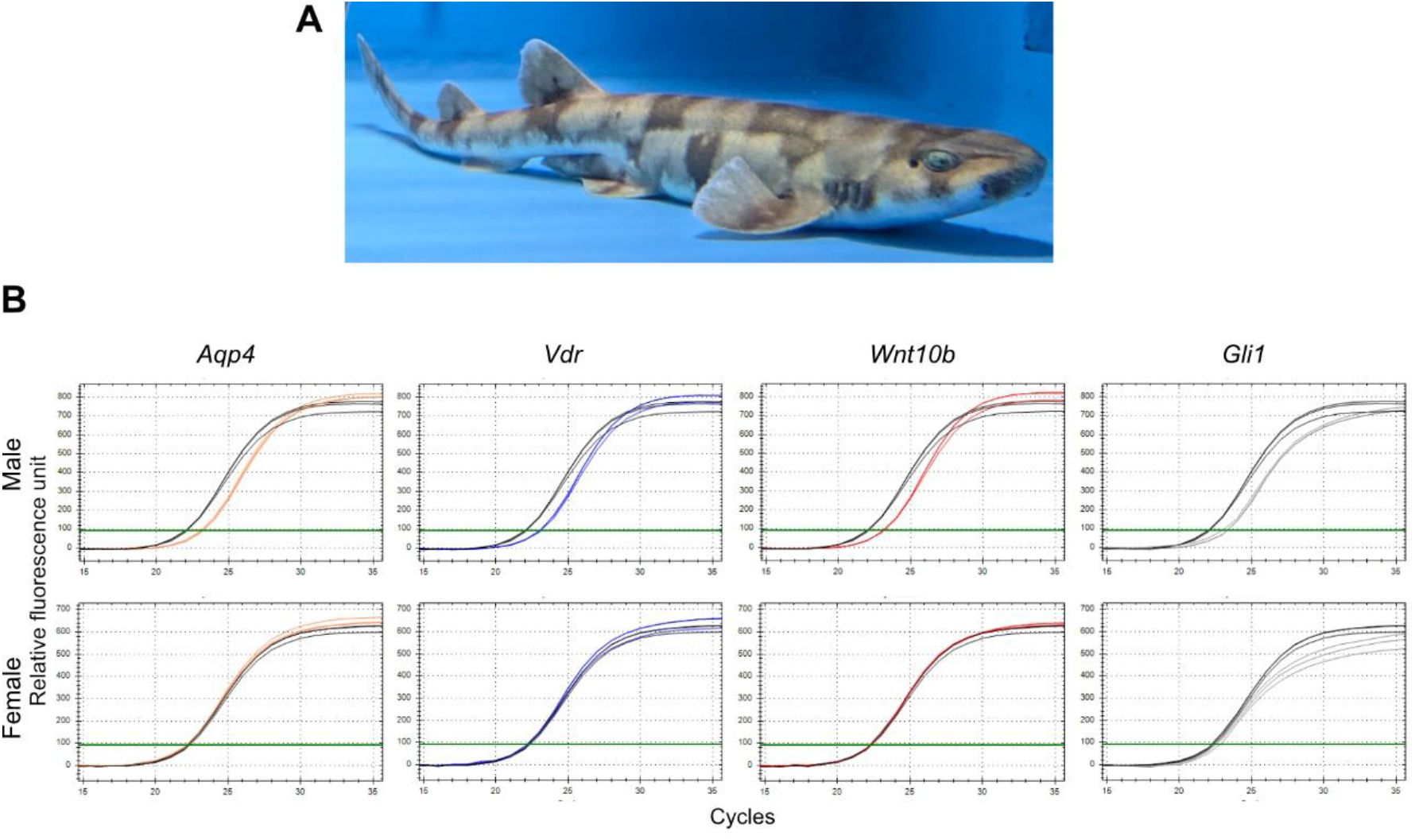
Sexing for cloudy catshark with quantitative PCR amplification profiles demonstrating X chromosome dosage differences. A, Adult cloudy catshark. Photo credit: Shigehiro Kuraku. B, Representative real-time PCR amplification plots for male (upper, m1 individual) and female (lower, f1 individual) genomic DNA samples in triplicates. Four X-linked target genes (*Aqp4, Vdr, Wnt10b*, and *Gli1*; Niwa et al., 2025) were amplified alongside an autosomal reference gene (*FoxG1*, black). See text for full names of these genes.

Using these primers, we performed a real-time PCR experiment in technical triplicates for three male and three female individuals, respectively, for which sex is unambiguously determined morphologically (see Methods; Fig. 4B). In males (XY), the amplification curves of the X-linked markers are delayed by approximately one PCR cycle (0.86 ± 0.04 for m1 and 1.08 ± 0.04 for m2) relative to the autosomal reference (Fig. 4B), reflecting a single copy of the X chromosome (Suppl. Table S2). In females (XX), the X-linked markers and the autosomal reference amplify simultaneously, indicating equal copy numbers (two copies each). This difference in cycle threshold (*Δ*Cq) enables the differentiation of sexes based on relative gene dosage.

## Discussion

In this study, we established a robust, fail-safe molecular sexing toolkit for three shark species through two distinct methodologies depending on the available genomic resources: multiplex end-point PCR targeting Y-linked markers, and quantitative PCR (qPCR) targeting X chromosome dosage. While Y-linked markers are intrinsically useful, conventional single-target “presence/absence” assays suffer from a critical vulnerability in forensic and archival contexts: the risk of false negatives. In degraded samples, such as dried fins or historical tissues, a simple PCR failure due to low DNA quality can easily be misinterpreted as a “female” result. To overcome this limitation, our framework inherently incorporates intrinsic internal controls. For species with identified Y-linked sequences (whale shark and brownbanded bamboo shark), we integrated an autosomal reference gene (*HoxA1*) into the multiplex PCR, where its consistent amplification guarantees DNA integrity. For species lacking known Y chromosomes (cloudy catshark), our novel quantitative approach intrinsically provides a relative copy-number control against an autosome (*FoxG1*). Importantly, these internal control genes, *HoxA1* and *FoxG1*, were previously subjected to careful gene repertoire surveys in elasmobranch genomes and found to exist as single copies (Hara et al., 2018). Both methodologies effectively eliminate the false-negative bias, providing the absolute reliability paramount for wildlife forensics, conservation management, and developmental biology.

The realization of this dosage-based sexing method is a direct corollary of recent breakthroughs in chondrichthyan genomics. Unlike teleost fishes, whose sex-determining genes have been extensively characterized for decades (Kitano et al., 2024), the sex chromosomes of cartilaginous fishes remained completely cryptic until very recently. It was the advent of chromosome-scale genome assemblies that finally unmasked their sex-linked genomic architectures (Sendell-Price et al., 2023; Yamaguchi et al., 2023; Niwa et al., 2025). This trajectory demonstrates how fundamental, discovery-driven genomics can be seamlessly translated into practical biological toolkits (van Oosterhout et al., 2025), complementing other DNA-based approaches targeting chondrichthyans (Kadota et al., 2023; Devloo-Delva et al., 2024; Bock et al., 2026).

Beyond conservation management, the new sexing solution for sharks and rays/skates offers a transformative advantage for developmental biology. In chondrichthyans, the molecular mechanisms underlying sex determination and gonadal differentiation remain poorly understood, representing a rapidly burgeoning frontier in vertebrate biology. A major bottleneck has been the inability to distinguish genetic sexes at early embryonic stages, long before the morphological manifestation of claspers. Our assay overcomes this hurdle by enabling definitive genetic sexing from tissue biopsies of early embryos (Niwa et al., 2025). By allowing researchers to confidently identify “future males” and “future females” prior to phenotypic sex differentiation, this method will facilitate further investigations into the developmental cascades that drive chondrichthyan sex determination.

Furthermore, the logistical challenges inherent to chondrichthyan research make this robust assay particularly valuable in “museomics” for historical ecology (Davis & Knapp, 2025). Because many shark species attain massive body sizes, the long-term archiving of intact whole specimens is spatially and financially prohibitive for most institutions. Consequently, researchers and museums frequently rely on spatially economical sample processing—storing small tissues, such as fin clips or blood samples, in freezers. These archival materials, much like the processed fins confiscated in the illegal wildlife trade, inevitably yield highly fragmented DNA. Conventional qualitative PCR frequently fails on such templates, risking false-female misidentifications. Our dosage-based qPCR approach, as well as our adoption of autosomal marker amplification, circumvents this limitation, ensuring that accurate demographic data and historical operational sex ratios can be reliably reconstructed even from these physically constrained, long-term archives.

Despite the robustness of our quantitative dosage-based and Y-targeted approaches, we acknowledge a current limitation, namely the unavailability of universal primers covering diverse elasmobranchs. In contrast to certain teleost lineages—such as salmonids, where the highly conserved *sdY* gene serves as a nearly universal, clade-wide sexing marker (Yano et al., 2013)—the shark Y chromosomes are divergent and remain largely uncharacterized across species (Niwa et al., 2025). However, as the repository of high-quality, chromosome-scale elasmobranch genomes continues to expand, future comparative studies may unveil master sex-determining genes possibly shared across broader taxonomic groups.

Looking forward, the expansion of this methodology to other heavily exploited species is a pressing priority. Pelagic species such as the blue shark *Prionace glauca*, silky shark *Carcharhinus falciformis*, and the shortfin mako *Isurus oxyrinchus* comprise a considerable proportion of the global fin trade and exhibit pronounced sexual segregation across different oceanic regions (Cardeñosa et al., 2020; Fields et al., 2018; Pacoureau et al., 2021). Y chromosome, or even the chromosome-level genome assemblies, has not yet been reported for the two latter species, but applying our robust sexing assay, demonstrated primarily for the cloudy catshark, to these species will facilitate sex-aware populational analysis.

While the present study relies on purified genomic DNA, the procedure could be substantially streamlined by adopting direct PCR from crude tissues, a technique routinely used in high-throughput genotyping of common laboratory species (Chum et al., 2012). Utilizing specialized DNA polymerases resistant to inhibitory contaminants would eliminate the DNA extraction step, offering a more rapid and field-friendly alternative for large-scale ecological surveys (Kermekchiev et al., 2009). Ultimately, integrating this DNA-based, sex-aware monitoring into routine workflows—spanning from embryonic research to global wildlife forensics—will provide the high-resolution data required to understand chondrichthyan biology and ensure the long-term persistence of their populations.

## Acknowledgments

The authors thank Mitsutaka Kadota for seminal ideas that led up to this project, Hatsune Makino-Itou and Akane Kawaguchi for advice in experimentation, animal caretakers including Itsuki Kiyatake at Osaka Aquarium Kaiyukan, Shinya Yamauchi at Aquamarine Fukushima, Tsuzuki Nobutaka at Shimoda Aquarium, and Rui Matsumoto at Okinawa Churaumi Aquarium, as well as Wataru Takagi and Susumu Hyodo at The University of Tokyo, for animal sampling in our earlier studies, and Chi Ju Yu for insightful discussion about potential application of the proposed methods. Computations were partially performed on the NIG supercomputer at the ROIS National Institute of Genetics.

## Funding

This research was supported by intramural grants from the National Institute of Genetics to SK. The funder played no role in study design, data collection, analysis and interpretation of data, or the writing of this manuscript.

## Author contributions

YU and SK conceived the study. AO and YK performed analyses. All authors contributed to final writing of the manuscript.

## Data availability

Nucleotide sequences of the genes used as PCR markers for the present study have been deposited in the public DNA sequence database.

---

**Supplementary Table 1.**
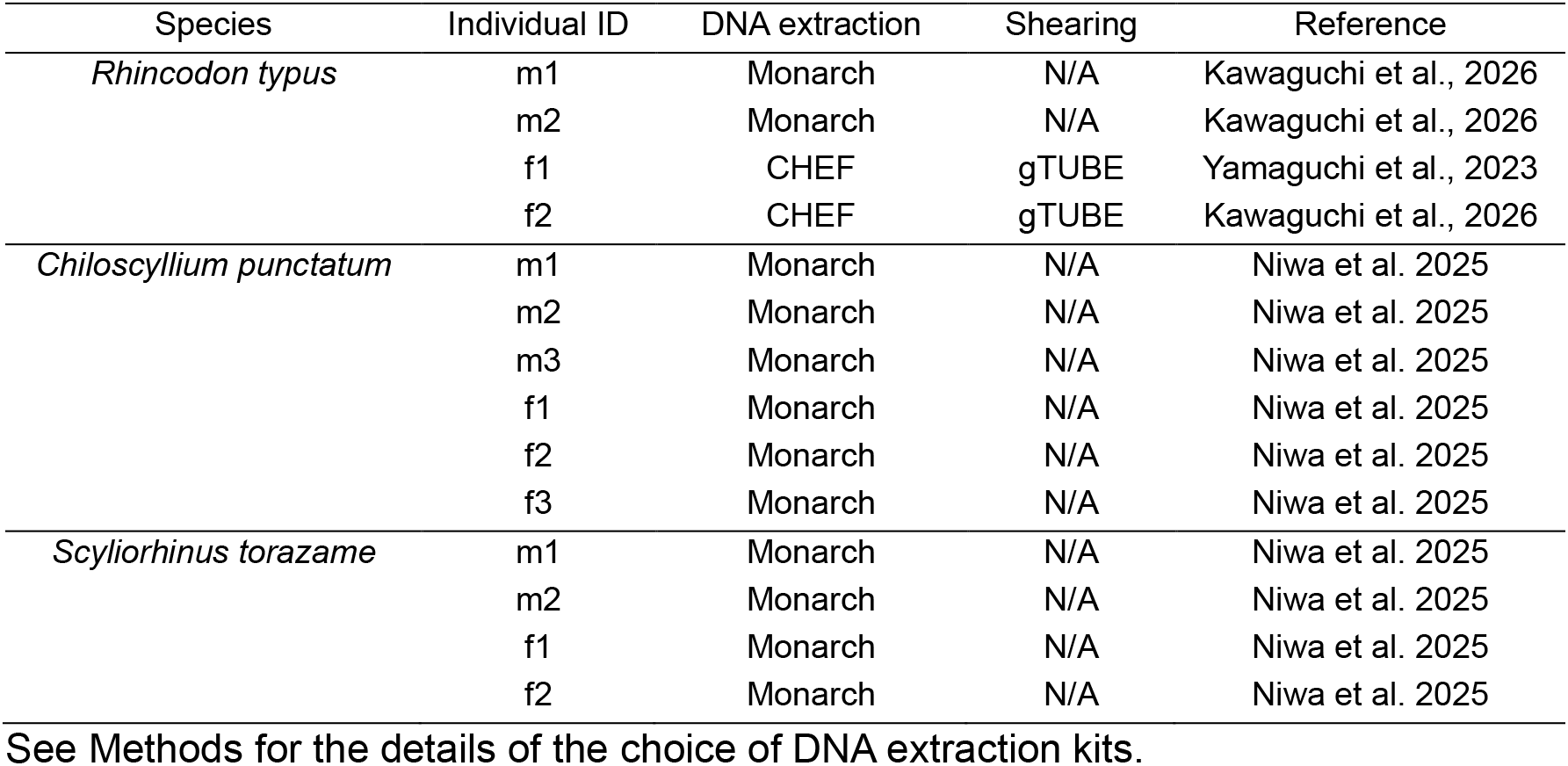
Genomic DNA samples used in this study.

| Species | Individual ID | DNA extraction | Shearing | Reference |
| --- | --- | --- | --- | --- |
| *Rhincodon typus* | m1 | Monarch | N/A | Kawaguchi et al., 2026 |
|  | m2 | Monarch | N/A | Kawaguchi et al., 2026 |
|  | f1 | CHEF | gTUBE | Yamaguchi et al., 2023 |
|  | f2 | CHEF | gTUBE | Kawaguchi et al., 2026 |
| *Chiloscyllium punctatum* | m1 | Monarch | N/A | Niwa et al. 2025 |
|  | m2 | Monarch | N/A | Niwa et al. 2025 |
|  | m3 | Monarch | N/A | Niwa et al. 2025 |
|  | f1 | Monarch | N/A | Niwa et al. 2025 |
|  | f2 | Monarch | N/A | Niwa et al. 2025 |
|  | f3 | Monarch | N/A | Niwa et al. 2025 |
| *Scyliorhinus torazame* | m1 | Monarch | N/A | Niwa et al. 2025 |
|  | m2 | Monarch | N/A | Niwa et al. 2025 |
|  | f1 | Monarch | N/A | Niwa et al. 2025 |
|  | f2 | Monarch | N/A | Niwa et al. 2025 |

## Notes

### Competing Interest Statement

The authors have declared no competing interest.

